# Placental gene expression mediates the interaction between obstetrical history and genetic risk for schizophrenia

**DOI:** 10.1101/147207

**Authors:** Gianluca Ursini, Giovanna Punzi, Qiang Chen, Stefano Marenco, Joshua F. Robinson, Annamaria Porcelli, Emily G. Hamilton, Marina Mitjans, Giancarlo Maddalena, Martin Begemann, Jan Seidel, Hidenaga Yanamori, Andrew E. Jaffe, Karen F. Berman, Michael F. Egan, Richard E. Straub, Carlo Colantuoni, Giuseppe Blasi, Ryota Hashimoto, Dan Rujescu, Hannelore Ehrenreich, Alessandro Bertolino, Daniel R. Weinberger

**Author notes:** Correspondence to: Daniel R. Weinberger, Lieber Institute for Brain Development, Johns Hopkins University School of Medicine, 855 North Wolfe Street, Baltimore, Maryland 21205.

## Abstract

Defining the environmental context in which genes enhance susceptibility can provide insight into the pathogenesis of complex disorders, like schizophrenia. Here we show that the intrauterine and perinatal environment modulates the association of schizophrenia with genomic risk, as measured with polygenic risk scores (PRS) based primarily on GWAS significant variants. Genomic risk interacts with intrauterine and perinatal complications (Early Life Complications, ELCs) in each of three independent samples from USA, Italy and Germany (overall n= 1693, p= 6e-05). In each sample, the liability of schizophrenia explained by PRS is nominally more than five times greater in the presence of a history of ELCs compared with its absence. In each sample, patients with positive ELC histories have higher PRS than patients without ELCs, which is further confirmed in two additional patient samples from Germany and Japan (overall n=2038, p= 1e-04). The gene set based on the schizophrenia loci interacting with ELCs is highly expressed in multiple placental compartments and dynamically regulated in placenta from complicated in comparison with normal pregnancies. The same genes are differentially up-regulated in placentae from male compared with female offspring. The interaction between genomic risk and ELCs is mainly driven by GWAS significant loci enriched for genes highly expressed in the various placenta samples. Molecular pathway analyses based on the genes not driving this interaction reflect previous analyses about schizophrenia risk-genes, while genes highly and differentially expressed in placentae implicate an orthogonal biology involving cellular stress. These results suggest that the most significant genetic variants detected by current schizophrenia GWAS contribute to risk in part by converging on a developmental trajectory sensitive to events affecting placentation, which may underlie the male preponderance of schizophrenia and offer new insights into primary prevention.

## INTRODUCTION

Schizophrenia is a complex disabling disorder occurring in all populations, with a lifetime morbid risk of around 0.7-0.8%^1^. The high heritability of the disorder indicates a major role for genetic variants in its etiology^2,3^; however, non-genetic influences involving the intrauterine environment have been repeatedly implicated in explaining at least part of the non-shared environmental contribution to the disorder^3-6^. Animal studies have shown that exposure to environmental insults *in utero* leads to altered response to stress postnatally, with effects on brain development and behavior that are partly mediated by gene expression changes in placenta^7-9^, a key environmentally sensitive organ during development^9,10^. Studies in animals also reveal that males are more vulnerable than females to prenatal adversities^8,9^.

An important role for the intrauterine environment in the etiology of schizophrenia is consistent with the disorder’s putative neurodevelopmental origins^11^ and is also supported by many epidemiological studies. For example, the prevalence of schizophrenia increases in offspring of mothers who were in the second trimester during influenza epidemics; in a prospective study, maternal respiratory infection during pregnancy increased the risk of schizophrenia in the offspring 3- to 7-fold^5,12^. More generally, schizophrenia has been associated with a number of early life complications (ELCs), i.e. potentially adverse events occurring during pregnancy and labor, at delivery, and early in neonatal life^5,12,13^. Meta-analyses of this body of literature have found that ELCs increase risk by 1.5- to 2-fold^13^, a greater effect than any common genetic variant. Studies of ELCs in high-risk individuals (i.e., offspring of parents affected with schizophrenia) suggest an interactive role for genetic background^13^, which is consistent with preliminary evidence of a relationship between ELCs, hypoxia-related genes, and risk of schizophrenia^13-15^.

Genome-wide association studies (GWAS) indicate that genetic risk of schizophrenia across heterogeneous samples is conferred by many small-effect alleles throughout the genome^16^. Studies of rare chromosomal defects showing greater penetrance also imply a myriad of susceptibility genes^17-19^, indicating that the genetic architecture of the disorder is heterogeneous, consistent with polygenic mechanisms^16,20^. While current GWAS are not designed to detect complex genetic and environmental heterogeneity^16^, we hypothesized that the most significant GWAS associations might achieve their statistical status by converging on early developmental mechanisms sensitive to environmental factors that are also relatively common among patients. Here, we analyze the role played by the intrauterine and perinatal environment in modulating the association of schizophrenia with genomic risk, with emphasis on the placental transcriptome.

## RESULTS

### Study Overview

We used a multi-step approach to investigate whether the intrauterine and perinatal environment modulates the association of schizophrenia with genomic risk and the role of the placenta. We first explored the interaction between genomic risk for schizophrenia and history of ELCs on case-control status in three, independent clinical samples (total n= 1693): a discovery sample of 501 individuals from USA (*scz_lie_eur*: 267 healthy subjects and 234 patients with schizophrenia, all Caucasian); an Italian sample of 273 subjects (*scz_bari_eur*: 182 healthy controls, 91 patients with schizophrenia, all Caucasian), and a German sample of 919 subjects (*scz_munc_eur*: 398 healthy subjects and 521 patients with schizophrenia, all Caucasian; see Table 1 and S1 for sample information). Genomic risk for schizophrenia was measured with polygenic risk score based on GWAS significant alleles (p<5e-08, PRS1; SNPs in Table S2)^16^, while ELCs history was assessed with the McNeil-Sjöström scale^21^, leading to a binary score (presence/absence of ELCs with severity level ≥4, see on-line methods and SI for details) for each ELC. We further tested the relationship between genomic risk for schizophrenia and ELCs score in two independent samples of only patients (total n=1192): another independent German sample of 1020 patients with schizophrenia (*scz_gras_eur*), and a Japanese sample of 172 patients with schizophrenia (*scz_osak_asi*) (Tables 1 and S1).

**Table 1.** Sample composition and summary statistics. Columns 1 and 3 report number (n) of healthy subjects (CONT) and patients with schizophrenia (SCZ) in the discovery sample (*scz_lie_eur*), in the 4 replication samples, and in the merged samples (*merged1*: cases and controls; *merged2*: cases only); columns 2 and 4 (“ELCs”) report frequency of ELCs in controls and patients with schizophrenia (NB: ELCs assessment was different among samples, while the scoring system was the same; see on-line methods and SI, “Considerations about the assessment and the frequency of ELCs”, for details). Columns 5 (“z”) and 6 (“p”) report the statistics of the association of ELCs with case-control status (NB: in *merged1* sample values within brackets refer to the statistics after adjustment for sample effect). Columns 7-15 report the statistics (“R2”: Nagelkerke R^2^ variance explained and “p”) for the relationship between PRS1 and case-control status in the whole sample (columns 8, 9), and in individuals without (11, 12) and with (14, 15) a history of ELCs. Columns 16-21 report the statistics for the relationship between PRS1 and ELCs binary scores in the whole sample (columns 16, 17), in controls (18, 19) and in patients with schizophrenia (20, 21). Columns 22-23 report the statistics for the interaction between PRS1 and ELCs on case-control status. NB: significant results (p<0.05) are highlighted in bold.

|  | CONT |  | SCZ |  | ELCs and diagnosis |  | PRS1 and diagnosis |  |  |  |  |  |  |  |  | PRS1 and ELCs |  |  |  |  |  | PRS1 inter |
| --- | --- | --- | --- | --- | --- | --- | --- | --- | --- | --- | --- | --- | --- | --- | --- | --- | --- | --- | --- | --- | --- | --- |
|  |  |  |  |  |  |  | whole sample |  |  | ELCs absence |  |  | ELCs presence |  |  | whole sample |  | CONT |  | SCZ |  |  |
|  | n | ELCs | n | ELCs | z | p | N | R2 | p | N | R2 | p | N | R2 | p | t | p | t | p | t | p | t |
| scz_lie_eur | 267 | 0.67 | 234 | 0.66 | -0.38 | 0.7 | 501 | 0.06 | 2e-7 | 167 | 0.01 | 0.28 | 334 | 0.11 | 5e-09 | 0.68 | 0.49 | -0.68 | 0.50 | 2.21 | 0.03 | 2.87 |
| scz_bari_eur | 182 | 0.51 | 91 | 0.47 | -0.51 | 0.61 | 273 | 0.02 | 0.04 | 138 | <.01 | 0.91 | 135 | 0.09 | 0.001 | 1.14 | 0.26 | -0.57 | 0.57 | 2.88 | 0.01 | 2.17 |
| scz_munc_eur | 398 | 0.15 | 521 | 0.24 | 3.54 | 4e-04 | 919 | 0.04 | 5e-8 | 733 | 0.02 | 1e-4 | 186 | 0.11 | 2e-05 | 0.91 | 0.36 | -1.64 | 0.10 | 1.60 | 0.05 | 2.12 |
| scz_osak_asi |  |  | 172 | 0.60 |  |  | 172 |  |  | 69 |  |  | 103 |  |  |  |  |  |  | 1.79 | 0.04 |  |
| scz_gras_eur |  |  | 1020 | 0.32 |  |  | 1020 |  |  | 690 |  |  | 330 |  |  |  |  |  |  | 1.70 | 0.04 |  |
| merged1 | 847 | 0.39 | 846 | 0.38 | -0.33<br>(1.97) | 0.74<br>(0.05) | 1693 | 0.04 | 1e-14 | 1038 | 0.02 | 2e-4 | 655 | 0.10 | 2e-15 | 1.49 | 0.14 | -1.58 | 0.10 | 3.25 | 0.001 | 4.02 |
| merged2 |  |  | 2038 | 0.37 |  |  | 2038 |  |  | 1281 |  |  | 757 |  |  |  |  |  |  | 3.86 | 1e-04 |  |

Second, we verified whether the genes mapping to the GWAS loci showing the strongest association with schizophrenia and interacting with ELCs are: i) enriched in placenta, relative to the expression of genes in the GWAS loci of lesser significance, ii) differentially expressed in placentae from complicated pregnancies compared with normal placental controls, iii) differentially expressed in placentae from male compared with female offspring. Finally, we evaluated the role of the GWAS SNPs marking loci containing genes highly expressed and dynamically modulated in placentae from complicated pregnancies in driving the interaction between PRS and ELCs; and, using the same set of placental enriched genes, we performed pathway, gene ontology and co-expression analyses.

### Interaction of polygenic risk profile score (PRS) and early life complications scores (ELCs) on case-control status

#### Discovery sample (scz lie eur)

In the USA discovery sample, the polygenic risk profile score constructed from alleles showing significant (p<5e-08) association with schizophrenia (PRS1), in a meta-analysis of PGC GWAS datasets after excluding the *scz_lie_eur* sample (SNPs in Table S2), was associated with case-control status (N=501, t= 5.347, p=2e-07) and accounted for approximately 6% of risk prediction (Nagelkerke’s pseudo*R*^2^ = 0.06; Fig. 1a and Table S3). In this sample, ELC scores were not significantly different among healthy subjects and patients with schizophrenia (z=-0.38, p=0.704). However, multiple logistic regression revealed a significant interaction between PRS1 and ELCs on case-control status (t=2.87, p=0.004; Fig. 1b); results of the multiple regression also indicated that the ELC score was associated with schizophrenia when covarying for genetic risk score (t=2.11, p=0.03), while PRS1 was not associated with schizophrenia when covarying for ELCs (t=1.18, p=0.24). We then analyzed the relationship between PRS1 and case-control status in the absence and in the presence of ELCs. This analysis revealed that the liability of schizophrenia explained by the genetic risk score was highly significant in the context of ELCs (N= 334, Nagelkerke’s pseudo*R*^2^ = 0.114, t=5.97, p=5.02e-09), but not in the absence of them (N= 167, Nagelkerke’s pseudo*R*^2^ = 0.008, t=1.07, p=0.28; Fig. 1b). We evaluated the same relationship in the context of each severity level of ELCs; strikingly, while in the absence of potentially serious ELCs (weights 0, 1, 2 and 3) cases and controls were not different in PRS1, the two groups became clearly differentiated as the severity of ELCs increased (Fig. 1c). These results were not affected by the inclusion or the exclusion of the top GWAS significant variant in the extended MHC region (chr6: 25-34 Mb; Table S4).

**Fig. 1.**
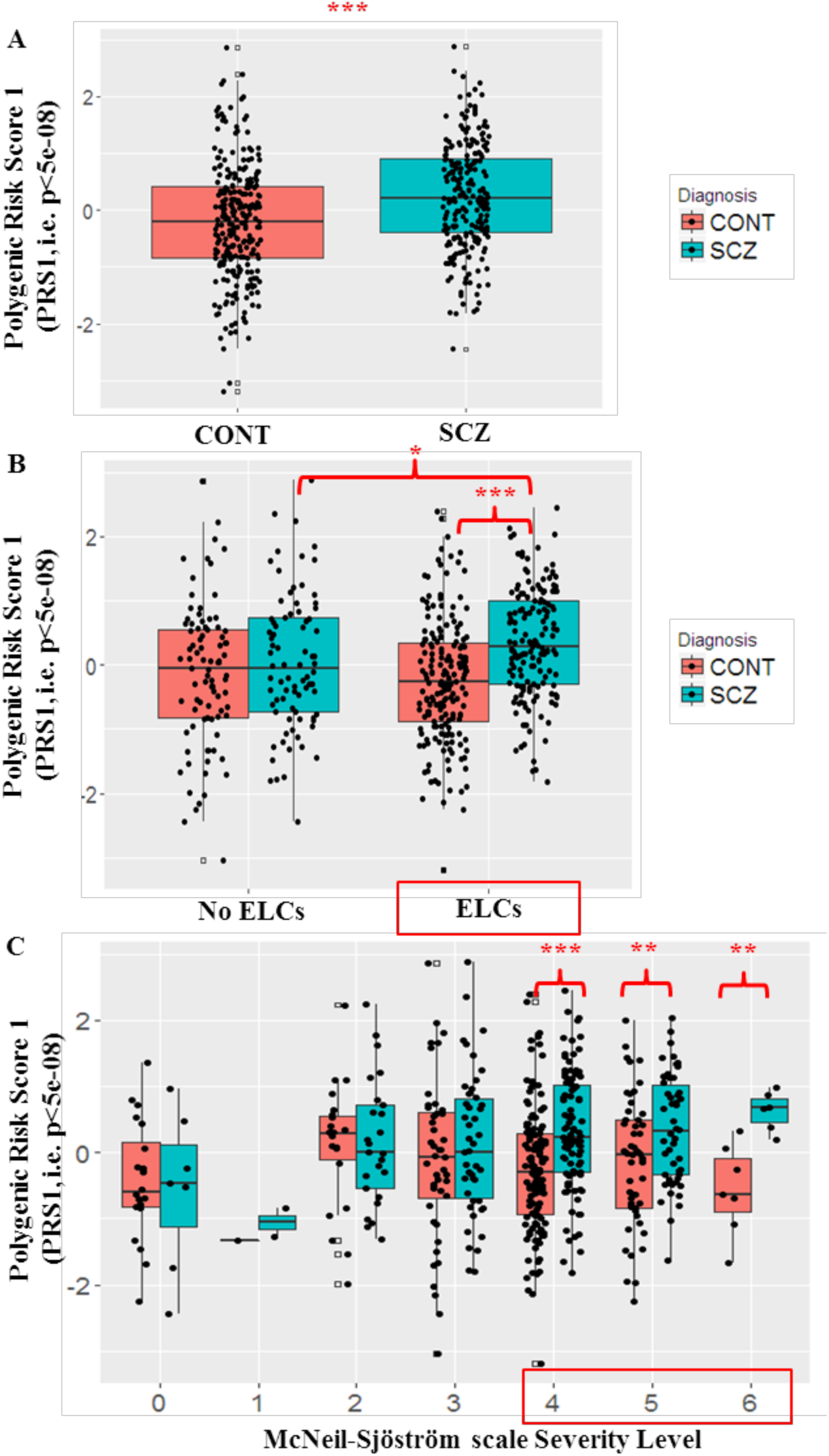
Polygenic Risk Score 1 (PRS1), Early Life Complications scores (ELCs) and schizophrenia. **A)** Boxplots of the relationship between PRS1, constructed from alleles showing significant association with schizophrenia (p<5e-08) and case-control status in the USA discovery sample (*scz_lie_eur)*: patients with schizophrenia (SCZ, blue box) have a greater PRS1 than controls (CONT, red box). **B)** Boxplots of the interaction between PRS1 and ELCs on case-control status: PRS1 and ELCs scores interact on case-control status, so that PRS1 is associated with schizophrenia principally in the context of ELCs (right) while not in absence of ELCs in this sample (left). **C)** Boxplots of the relationship between PRS1 and case-control status in the context of ELCs with different severity levels: patients with schizophrenia (SCZ, blue box) have greater PRS1 than controls (CONT, red box), only when exposed to ELCs potentially clearly harmful for the offspring CNS; the difference is particularly evident in the presence of ELCs associated with “very great harm or deviation in offspring” (severity level 6). NB: *: p<0.05; **: p<0.01; ***: p<0.001; ELCs with severity scores ≥ 4 are considered harmful or relevant factors in fetal stress, while ELCs with severity scores 0,1,2,3 are not.

To further represent the capacity of PRS1 to predict case-control status in the context of ELCs, we grouped individuals in PRS1 quintiles and estimated odds ratios (ORs) for each quintile with reference to the lower risk quintile, stratifying by ELCs status, using a standard epidemiological approach to a continuous risk factor^16^. The OR increased with the PRS1 quintiles only in the sample with ELCs, maximizing in the highest quintile, where it reached an OR of 7.99 (95% confidence interval [CI]: 3.6-17.8, p=1.19e-07) in the presence of ELCs, and only 1.73 (CI: 0.66-4.55, p=0.26) in the absence of ELCs (Fig. 2a).

**Fig. 2.**
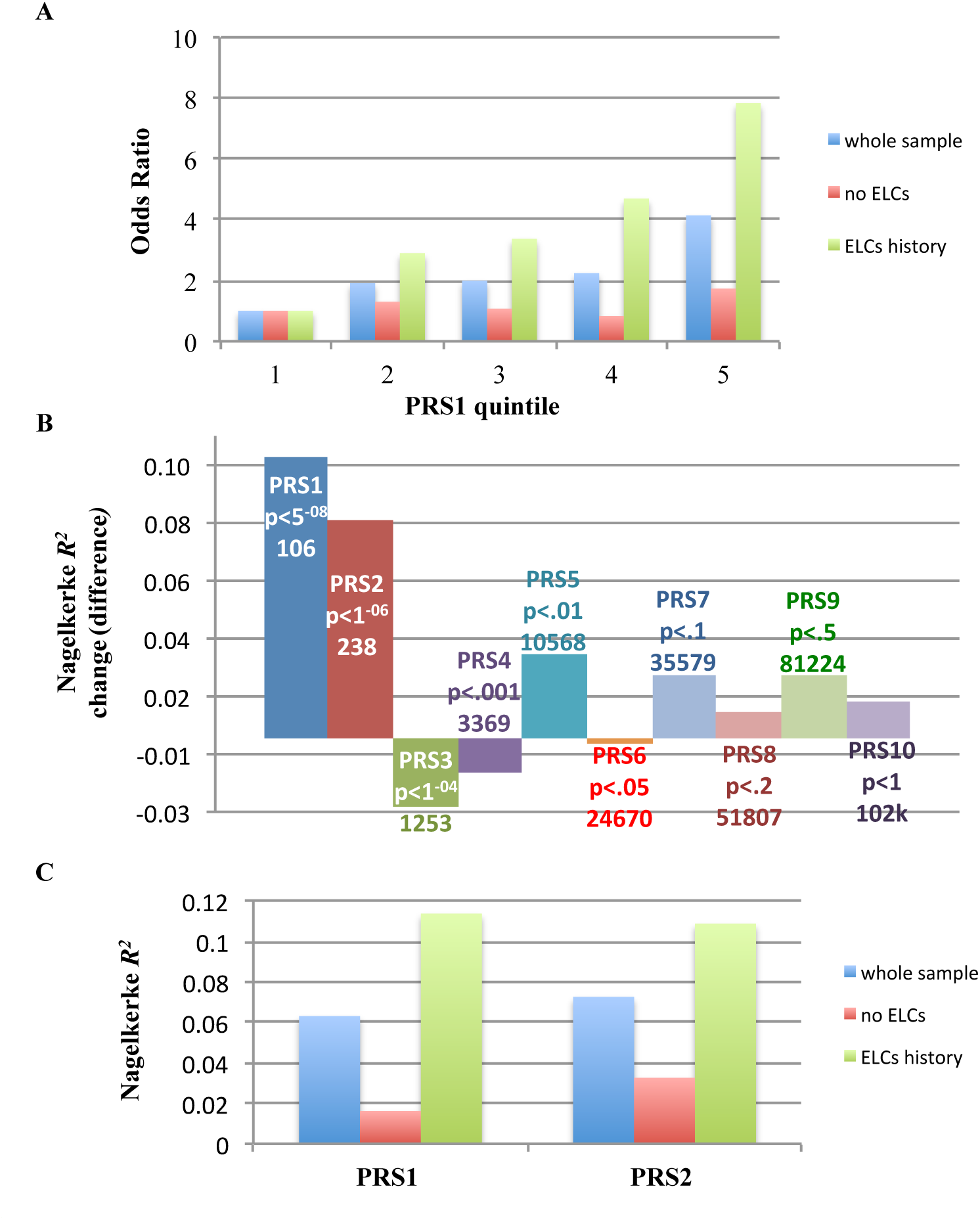
History of ELCs affects the variance explained by the most significant PRS’s (i.e. PRS1 and PRS2) in the USA discovery sample (*scz_lie_eur)*. **A)** Odds ratio (OR) for schizophrenia by PRS1 and ELCs history: the PRS constructed from alleles showing significant association with schizophrenia (p<5e-08) was converted to quintiles (1=lowest PRS, 5=highest PRS) and the ORs were estimated for affected status for each quintile with reference to the lowest risk quintile, in the whole sample (blue bars) and in absence (red: ELCs score = 0) and presence (green: ELCs score= 1) of ELCs; the OR strongly increases with the PRS1 in the context of a history of ELCs, while not in the absence of ELCs. **B)** Change of variance explained by PRS’s in presence of ELCs compared to the absence of ELCs: proportion of variance (Nagelkerke R^2^) of schizophrenia risk was calculated in presence and absence of ELCs history for each of the 10 different PRS’s constructed from alleles showing association with schizophrenia at different threshold of significance (the data labels of each bar report the corresponding PRS, the GWAS p-value of association with schizophrenia of the SNPs included in each PRS, and the number of SNPs in each PRS). Shown is the difference (y-axis) between Nagelkerke R^2^ in presence of ELCs and the Nagelkerke R^2^ in absence of ELCs history. The change in variance explained based on ELCs history gradually decreases when considering different PRS’s constructed from alleles showing association with schizophrenia at lower thresholds of significance (i.e., not GWAS-significant), so that only PRS1 and PRS2 interact with ELCs (NB: a difference higher than 0 indicates greater proportion of variance explained by the PRS in presence of ELCs than in absence). **C) Proportion of variance of schizophrenia by PRS1 and PRS2, and ELCs history:** Nagelkerke R^2^ (y-axis) of PRS1 and PRS2 are greater in the presence of ELCs (green bars: ELCs score= 1) than in the absence of ELCs (red bars: ELCs score= 0), suggesting that the aggregate effect of the GWAS-significant SNPs (PRS1) and the almost GWAS-significant SNPs (PRS2) is conditioned on ELCs history, differently from the other PRS’s **(B)**.

We then analyzed whether the interaction between genomic risk and ELCs was specific for the PRS constructed with GWAS-significant alleles (PRS1) or was also found with other PRS levels (i.e. PRS2 to 10) constructed from alleles showing association with schizophrenia at lesser statistical thresholds (i.e. not GWAS significant). Interestingly, the ELCs-dependent change in schizophrenia risk variance, explained by PRS, gradually decreased when considering different PRS’s constructed from variants showing association with schizophrenia at the lower thresholds of significance (Fig. 2b). Specifically, only the first two scores, constructed from the alleles showing the strongest clinical association with schizophrenia (PRS1=p<5e-08; and PRS2=p<1e-06), interact with ELCs on case-control status, and the variance in risk explained by them is much higher in individuals with a history of ELCs, compared with those without (Fig. 2b-c and Fig. S1-2). The other scores, PRS3to10, do not show any interaction and the variance explained by them is not influenced by a history of ELCs (Fig.2b and S1-2). This is consistent with the possibility that the latter scores - involving a greater number of putative susceptibility genes - capture a greater number of genetic risk variants acting in a simply cumulative way, while the aggregate effect of the GWAS-significant SNPs (PRS1) and the almost GWAS-significant SNPs (PRS2) is more conditioned on the history of ELCs (Fig. 2c and S1-2). These results strongly suggest that the reason PRS1 and PRS2 loci achieve their privileged statistical significance status in this heterogeneous clinical sample is because of this interaction. From another perspective, the data show that patients with a history of ELCs have greater PRS1 than patients without ELCs (N= 234, t=2.21, p=0.028), whereas this relationship is not seen in healthy subjects (N= 267, t=- 0.68, p=0.50). Maternal PRS’s were available on a subsample of healthy mothers (N=160) of schizophrenic offspring and were also not associated with ELCs in their offspring (t= 0.08, p=0.93, Table S5).

#### Replication samples

##### Italian sample (scz_ bari_eur)

As in the discovery sample, PRS1 was positively associated with case-control status (N=273, *R*^2^=0.02, t=2.11, p=0.036, Table S3) in the Italian replication sample. ELC scores were not differentially distributed between healthy subjects and patients with schizophrenia (z=-0.51, p=0.61). Even in this smaller sample, PRS again showed a significant interaction with ELCs in predicting case-control status (t=2.17, p*_one-sided_*=0.015; Table 1 and Fig. S3a). When analyzing the relationship between PRS1 and case-control status in the context of ELCs, this PRS was once again associated with schizophrenia only in the presence of ELCs (N=135, t= 3.38, p=0.001), and not in their absence (N= 138, t= -0.11, p=0.91), so that the variance of case-control status explained by PRS1 was much higher in individuals with a history of ELCs (*R*^2^=0.09, Fig. S3b) than in those without such history (*R*^2^=0.0001, Fig. S3b); and again subjects who experienced ELCs who were in the upper quintile for PRS1 had the highest OR (7.33, 95% CI: 1.8-29.8, p =0.003). This replication analysis also confirmed that the PRS was associated with ELCs only in cases (N= 91, t=2.88, p*_one-sided_*=0.005), but not in controls (N= 182, t=-0.57, p=0.57, Table 1).

##### German sample 1 (scz_munc_eur)

We next analyzed the interaction between PRS and ELCs in a larger sample from Germany, where the history of ELCs was assessed using a questionnaire based on only a few items, thus increasing the chance of incomplete information. As expected, the frequency of ELCs (0.20) was lower in this sample, compared with the others, given the lower sensitivity of the questionnaire for ELCs assessment (see SI for details). Once again, PRS1 was positively associated with case-control status (N=919, R^2^= 0.04, t=5.51, p=5e-08; Table S3). In this larger sample, we did find a significant association of ELC history with schizophrenia (z=3.54, p=0.0004, OR=1.85, 95% CI: 1.32-2.61). Despite the differences in ELC assessment compared with the earlier samples, we again found a significant interaction between PRS1 and ELCs on case-control status (t=2.12, p*_one-sided_*=0.017: Fig. S3c). More dramatically, and as in the other two samples, the variance of schizophrenia risk explained by PRS1 was nominally five times greater in the presence of ELCs (N= 186, *R*^2^ = 0.11; OR= 15.13, 95% CI: 4.32-52.98, t= 4.45, p=2e-05, Fig. S3d), than in their absence (N= 733, *R*^2^= 0.02; OR= 2.11, 95% CI: 1.33-3.37, t=3.88, p= 0.0001, Fig. S3d), although in this sample, the association of PRS1 and diagnosis was significant in both contrasts (Table 1). Consistent also with the other two samples, we found a statistical trend for a positive relationship between PRS1 and ELCs in patients with schizophrenia (N=521, t=1.60, p*_one-sided_*=0.0547), but not in controls (N= 398, t= -1.64, p=0.10), where it again tended towards the opposite direction.

##### Japanese sample (scz_ osak_asi)

In the Japanese sample of patients with schizophrenia, we tested whether cases with a history of ELCs had higher PRS’s than cases with no history of ELCs. It should be noted that the PRS derived from the European Caucasian sample of the recent GWAS study of schizophrenia has much less association with schizophrenia in Asian samples^16^, as would be expected because of genomic heterogeneity and different LD structure. However, since many of the alleles comprising the score likely monitor ancient haplotypes, an association with ELCs might still be expected. As in the other samples, we again found that PRS1 was associated with ELCs (N= 172, t=1.79, p*_one-sided_*= 0.047; Fig. S4), so that patients with a history of complications had higher PRS than patients who did not experience ELCs.

##### German sample 2 (scz_gras_eur)

We analyzed the relationship between PRS and ELCs in another independent sample of patients with schizophrenia from Germany, namely the Göttingen Research for Schizophrenia (GRAS) data collection^22,23^. Again, we found a positive association between PRS and ELCs, so that PRS1 was higher in patients with a history of ELCs, compared with patients without ELCs (N= 1020, t= 1.70, p*_one-sided_*= 0.044; Fig. S4).

##### Merged samples

We performed additional analyses in merged samples of cases and controls (*scz_lie_eur, scz_bari_eur, scz_munc_eur*) and of only cases (*scz_lie_eur, scz_bari_eur, scz_munc_eur, scz_osak_asi, scz_gras_eur*), after normalization of PRS’s in each population. In these merged samples, we confirmed the interaction of PRS1 and ELCs on case-control status (N= 1693, t= 4.02, p= 6.18e-05; Fig. S5) and the relationship between PRS and ELCs in patients with schizophrenia (N=2038, t= 3.86, p=0.0001). Also in the merged samples, only PRS1 and PRS2 interact with ELCs on case-control status, and only PRS1 and PRS2 are positively associated with ELCs in patients with schizophrenia (Fig. S6). The positive association between genomic risk and ELCs was not present in controls, where the trend was actually negative (p=0.10, Table 1), compatible with a pattern of a gene-environment interaction. Sensitivity analyses with sex and age as covariates and related interaction terms (in addition to genetic principal components) confirmed the same results (Table S6-7). These consistent results in five independent samples support the hypothesis that these top PRS’s are relevant for the etiopathogenesis of schizophrenia particularly in the context of ELCs, while other PRS’s (i.e. PRS3to10) may capture polygenic mechanisms of schizophrenia not directly related to ELCs.

### Expression of schizophrenia risk associated genes in placenta

While several recent studies show preferential expression of many schizophrenia risk genes in fetal brain^24-26^, the relationship between the PRS and ELCs that we found in five independent clinical samples from diverse ancestries points to the intrauterine context as a likely place where some risk genes for schizophrenia and environmental adversity intersect, with implications not limited to the brain. We therefore tested whether the genes mapping to the loci showing the strongest association with schizophrenia and interacting with ELCs (i.e., PRS1 and PRS2 genes; Fig. 2b) are more highly expressed in placenta, compared with randomly selected genes contributing to PRS’s constructed from alleles showing association with schizophrenia at lesser thresholds of significance (p>1e-06), which also do not show an interaction with ELCs (i.e. PRS3 to 10). We analyzed RNA sequencing data from placental tissue, generated in the Epigenome Roadmap Project (GSE16368), and found relatively greater expression of the PRS1 and PRS2 genes (N=1643 genes), compared with same size set of genes randomly selected from PRS3to10 genes (N=18029 genes), in multiple placental tissue compartments: amnion (n=4 samples, p=1e-04), basal plate (n=4, p=1e-04), chorion (n=4, p=3e-04), villi (n=4, p=1e-05), trophoblast (n=4, p=1e-05; 2^nd^ trimester: n=2, p=3e-05; 3^rd^ trimester: n=2, p=1.6e-06; Table S8). These results indicate that, as predicted, genes mapping to GWAS significant loci that interact with ELCs are more abundantly expressed in placenta than are genes in the other GWAS loci, which do not interact with ELCs.

### Differential expression of schizophrenia risk associated genes in placentae from complicated pregnancies

To elaborate on a specific role for the placenta in mediating the interaction between schizophrenia risk-genes and ELCs, we next analyzed whether the PRS1 and PRS2 genes were differentially expressed in placentae from pregnancies complicated with preeclampsia and/or intrauterine growth restriction (IUGR) compared with normal placental controls. Preeclampsia and IUGR are classic dangerous ELCs (severity level ≥ 4 in the McNeil-Sjöström scale) that have been associated – among other ELCs – with increased risk for schizophrenia^13,27,28^, and also where the primary affected cells have been isolated and studied *ex vivo*. In analyzing eight publicly available datasets, we consistently detected enrichment of the PRS1 and PRS2 genes (Table 2 and Table S9, see also Supplementary Information) among the genes differentially expressed in whole placentae from pregnancies complicated with preeclampsia (GSE24129: p=3.5e-04; GSE35574: 0.04; GSE10588: 0.003; GSE25906: 0.02) and IUGR (GSE24129: p=0.019; GSE35574: 0.007; GSE12216: 0.01), in preeclamptic cytotrophoblast (GSE40182: p=0.009) and chorionic villi (GSE12767: p=0.003), and in microvascular endothelium from IUGR/preeclamptic pregnancies (GSE25861: p=0.006). Since the PRS1 and PRS2 genes were among the highly expressed placental genes, we performed a sensitivity analysis controlling for average gene expression level and the results are consistent (Table S10). We observe that PRS1 and PRS2 genes tend to be up-regulated (positive t-statistics) in multiple placental samples from preeclampsia and IUGR, compared with placental controls.

**Table 2.** Differential expression of schizophrenia risk genes in placentae from complicated pregnancies. Genes mapping to loci showing the strongest association with schizophrenia (p<5e-08: PRS1, and p<1e-06: PRS2) and interacting with ELCs were tested for enrichment among the differentially expressed genes in preeclamptic and IUGR placental samples compared with controls, and in non-invasive CTBs, in 9 independent datasets (11 differential expression analyses: rows 1-11). The table shows the results on the gene set test analysis using the moderated t-statistics from each differential expression analysis (see SI for details): the PRS1 and PRS2 genes tend to be more highly ranked in differential expression compared to randomly selected genes from the other GWAS loci (PRS3to10). The χ2 test results confirm that PRS1 and PRS2 genes are enriched for differentially expressed genes compared with the remaining genes. Similar analyses in datasets from liver, stomach, heart, and from cells of embryonic origins, do not show any significant enrichment (last 10 rows), supporting the specificity of differential expression of PRS1 and PRS2 genes in complicated placentae.

| Dataset | Condition | Tissue | p-value of Gene-Set Test | $\chi^2$ test | |
| --- | --- | --- | --- | --- | --- |
| | | | | p-value | $\chi^2$ |
| GSE24129 | Preeclampsia | whole placentae | 3.5e-04*** | 0.002** | 7.93 |
| GSE24129 | IUGR | whole placentae | 0.019* | 0.0007*** | 10.21 |
| GSE35574 | Preeclampsia | whole placentae | 0.04* | 0.062 | 2.36 |
| GSE35574 | IUGR | whole placentae | 0.007** | 0.03* | 3.56 |
| GSE10588 | Preeclampsia | whole placentae | 0.003** | 0.002* | 8.46 |
| GSE25906 | Preeclampsia | whole placentae | 0.02* | 0.03* | 3.40 |
| GSE12216 | Preeclampsia | whole placentae | 0.01* | 0.01* | 4.82 |
| GSE40182 | Preeclampsia | CTB | 0.009** | 0.04* | 3.04 |
| GSE12767 | Preeclampsia | 1<sup>st</sup> trimester chorionic villi | 0.003** | 0.005** | 6.78 |
| GSE25861 | Preeclampsia/IUGR | microvascular endothelium | 0.006** | 0.04* | 3.002 |
| GSE65271 | CTB invasiveness | CTB | 0.005** | 0.002** | 8.30 |
| GSE28619 | Hepatitis (liver) | liver | 0.136 | 0.10 | 1.70 |
| GSE41804 | Hepatitis (liver) | liver | 0.285 | 0.20(opposite direction) | 0.69 |
| GSE27411 | HP gastritis (corpus) | stomach corpus | 0.377 | 0.45 | 0.01 |
| GSE27411 | HP gastritis (antrum) | stomach antrum | 0.453 | 0.34 | 0.17 |
| GSE42955 | Dilatative cardiomyopathy | heart | 0.172 | 0.40 | 0.07 |
| GSE3586 | Dilatative cardiomyopathy | heart | 0.283 | 0.10 (opposite direction) | 1.63 |
| GSE4172 | Dilatative cardiomyopathy | heart | 0.246 | 0.42 | 0.04 |
| GSE4483 | Hypoxia | 2<sup>nd</sup> trimester astrocytes | 0.470 | 0.18 | 0.85 |
| GSE26420 | MIBP1 overexpression | HEK293 cells | 0.263 | 0.001(opposite direction) | 8.01 |
| GSE64699 | IUGR | adipocytes from UC-MSC lines | 0.109 | 0.37 | 0.12 |

We considered the possibility that differential expression of PRS1 and PRS2 genes in complicated placentae might be a nonspecific response to pathology in adult or fetal tissue. We therefore performed the same differential expression analyses on datasets from tissues with diseases likely unrelated to schizophrenia, such as hepatitis (GSE28619, GSE41804), Helicobacter Pylori gastritis infection (GSE27411, with separate analysis in stomach corpus and antrum), and dilatative cardiomyopathy (GSE42955, GSE3586, GSE4172), as well in datasets from embryonal cells (GSE4483: astrocytes from 2^nd^ trimester fetal brain exposed to hypoxia/normoxia; GSE26420: HEK293 with a MIBP1 overexpression/control comparison, GSE64699: adipocytes differentiated from primary UC-MSC lines isolated from SGA and normal neonates). The PRS1 and PRS2 genes were not enriched among the genes differentially expressed in the pathological compared with normal condition in any of these datasets, from adult tissues and embryonic cells (Tables 2 and S9, see SI for details).

Since both preeclampsia and IUGR are ELCs attributed to a shallow invasion of the uterine interstitial compartment from cytotrophoblasts,^13,27-29^ we tested in a further dataset (GSE65271) whether the PRS1 and PRS2 genes are enriched among the genes differentially expressed in cytotrophoblasts cultured *ex vivo* under conditions of different invasiveness. Consistently, we found that PRS1 and PRS2 genes are up-regulated in low-invasive compared with high-invasive cytotrophoblasts (CTBs) cultured under hypoxic conditions (GSE65271: p=0.005; Table 2). Taken together, these results converge on the conclusion that schizophrenia GWAS-significant risk-associated genes that interact with ELCs are highly expressed in the placenta during early life and dynamically modulated in the placenta during biological stress, as reflected in their differential expression in placentae from complicated pregnancies, and that these associations are relatively placental specific.

### The genes highly expressed and dynamically modulated in placenta drive the interaction between PRS and ELCs on schizophrenia risk

The enrichment of expression in the placenta of genes in schizophrenia GWAS-significant loci provides circumstantial evidence that the interaction of these loci with ELCs on risk for schizophrenia arises at least in part because of primary effects in the placenta. To achieve a more direct test of this possibility, we created new PRS’s based on the GWAS SNPs marking loci containing genes highly expressed in normal placentae and dynamically modulated in placentae from complicated pregnancies (Table 2, and Table S10) and compared their interaction with ELCs to PRS’s derived from the SNPs marking the remaining GWAS significant loci first in our discovery sample (*scz_lie_eur*, Fig. 3a-d). The PRS’s from the former group significantly interact with ELCs in increasing risk for schizophrenia (PlacPRS1 [PRS1 ‘placental’ subset based on 56 SNPs]: t=2.86, p=0.004; PlacPRS2 [PRS2 ‘placental’ subset based on 112 SNPs]: t=3.10, p=0.002; Fig. 3a,c), while those from the latter do not (NonPlacPRS1 [PRS1 ‘non-placental’ subset based on 49 SNPs]: t=0.78, p=0.43; NonPlacPRS2 [PRS2 ‘non-placental’ subset based on 125 SNPs]: t= -0.53, p=0.60; Fig. 3b,d). Similar results were found in the other case-control samples (Fig. S7 and S8). Analyses on the merged samples of cases and controls (*scz_lie_eur*, *scz_bari_eur*, *scz_munc_eur*: n=1693, Fig. S9a-d) confirm these results; PlacPRS1 (t= 3.19, p= 0.0014: Fig. S9a) and PlacPRS2 (t= 3.28, p=0.0011: Fig. S9c) significantly interact with ELCs on case-control status, while nonPlacPRS1 and nonPlacPRS2 do not (Fig. S9b,d).

**Fig. 3.**
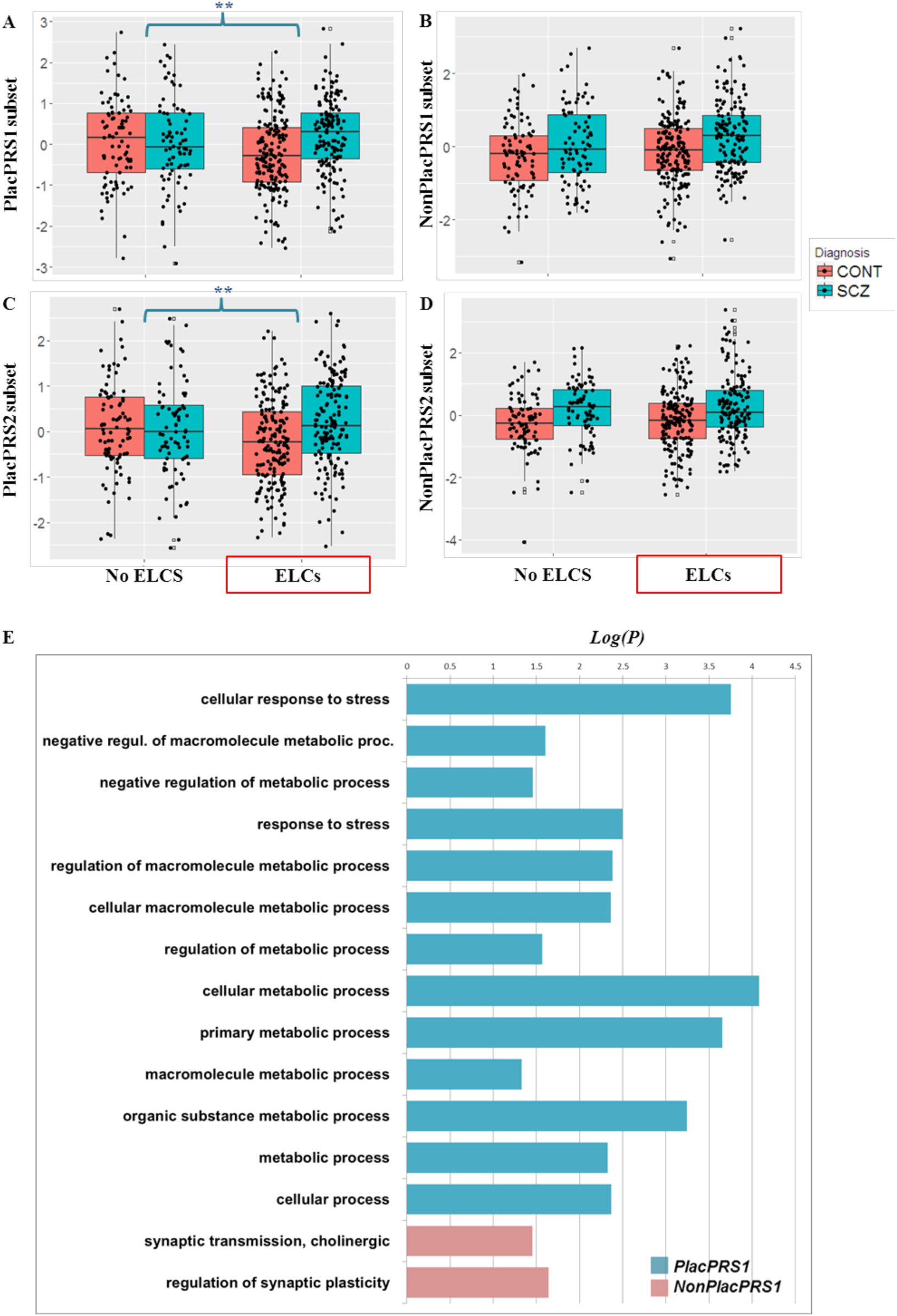
Schizophrenia risk genes dynamically modulated in placenta drive the interaction between genetic risk and ELCs. **A-D)** Using GWAS SNPs marking loci containing genes highly and differentially expressed in preeclamptic/IUGR placental samples, we created new PRSs (PlacPRS’s) and compared their interaction with ELCs to PRS’s derived from the SNPs marking the remaining GWAS significant loci (NonPlacPRS’s). The figure shows the boxplots of the interaction between PRS’s and ELCs on case-control status in the USA discovery sample (*scz_lie_eur)*: while PlacPRS’s significantly interact with ELCs on case-control status (**A and C**), NonPlacPRS’s constructed from the remaining loci do not (**B and D**). Similar results are found in the other clinical samples (Fig. S7-9). NB: **:p<0.01; the asterisks refer to the significance of the interaction. **E)** Biological processes GO terms enriched for PlacPRS1 and NonPlacPRS1 genes: PlacPRS genes implicate biological processes related to the fetal/placental response to stress, while NonPlacPRS genes implicate an orthogonal biology related to synaptic function (see also Supplementary Information, Tables S14-17, and Figures S10-S20).

To verify the specificity of these interactions to placenta gene expression, we calculated PRS’s based on the genes highly expressed in various adult and fetal tissues/embryonic cells, and differentially expressed in these organs during pathological/stress compared with the normal condition, employing the same procedure that we used for the computation of PlacPRS’s and NonPlacPRS’s (see SI for details). We also calculated brain PRS’s, based on SNPs in PRS1 and PRS2 loci associated with methylation quantitative trait loci (meQTL) in adult brain^30^ and with chromatin interaction in fetal brain^24^. In all of these sensitivity analyses, the PRS’s comprising SNPs marking loci having genes highly expressed in these diverse adult and fetal tissues and dynamically regulated in pathology of heart, liver and stomach, and in pathological cells of embryonic origins, do not significantly interact with ELCs on risk for schizophrenia (all p> 0.16 after FDR-correction, results in Tables S11-13), while only the SNPs mapping to the loci highly expressed and differentially expressed in placenta do (see SI for details).

### Biological insights about placental enriched genes associated with ELCs

Both PlacPRS1 and PlacPRS2 genes are significantly enriched for many pathways related to metabolic and cellular stress and hypoxia, particularly to “unfolded protein response”, “mitochondrial dysfunction” and “HIF1α signaling” (Fig. S10-11; Table S14), while not a single significant pathway enrichment could be obtained from the remaining PRS1 and PRS2 genes (NonPlacPRS1 and NonPlacPRS2), as well as from the whole PRS1 and PRS2 gene-sets, consistent with the analogously negative results of the original analysis of the GWAS significant loci^16^ (details in SI). It is noteworthy that the pathways (Fig. S10-11; Table S14), biological function and processes (Fig. 3e and S12-14, Tables S15-16), and cellular compartments (Fig. S15-S16, Table S16) implicated in PlacPRS genes are virtually orthogonal to those highlighted in other analyses of schizophrenia loci, such as synaptic function, calcium signaling, Fragile X associated proteins and chromatin remodeling^16^. Interestingly and in contrast, genes in the NonPlacPRS’s do tend to implicate some of these brain relevant functions. These results suggest that the loci containing the schizophrenia associated genes dynamically modulated and most enriched in the placenta contribute to schizophrenia risk at least in part by influencing the fetal/placental response to stress (Fig. S17-19), as exemplified by the cellular stress response gene HSF1^31,32^ being the main transcriptional regulator of genes in PlacPRS2 (Fig. S17-18; Table S17; see SI for details).

Because maternal stress during pregnancy in animals^8^ has been associated with inflammatory responses in the placenta, we performed an exploratory co-expression analysis of over 500 genes related to inflammation and the immune response in the placenta datasets from complicated pregnancies (see supplementary methods and Table S18-19). The PlacPRS1 and PlacPRS2 genes were significantly co-expressed with immune response genes, and in contrast to NonPlacPRS1 and NonPlacPRS2 genes as well as similarly sized gene sets of non-schizophrenia associated genes in the same datasets (Table S19 and Fig. S20). Interestingly, the PlacPRS1 and PlacPRS2 genes were also significantly co-expressed with “heat shock proteins” and “complement”genes (Table S19).

The suggestion of a distinct and orthogonal biology for the placental component of genomic risk raises the question of whether genetic prediction might be enhanced by deconstructing genomic risk into discrete sub-compartments that represent alternate risk biologies. We performed an exploratory analysis of this possibility. Previous studies have shown, as confirmed in our sample, that the prediction accuracy of PRS’s increases to an asymptote from PRS1 to PRS6. Since our data show that PRS1 interacts with ELCs, differentially when compared with PRS3to10, we excluded the SNPs in PRS1 from the calculation of the other PRS’s. We found that the aggregate effect on prediction accuracy of the SNPs contributing to PRS3to10 is higher when separating the contribution of PRS1 (Fig. S21). This is particularly true in the context of a history of ELCs, for each PRS. These results suggest that decomposing PRS’s based on early environmental exposure and placental genetic risk may increase the prediction accuracy of genetic variation for schizophrenia.

### Sex-specific analyses

The interaction between ELCs and genetic risk for schizophrenia is consistent with a body of literature pointing to the placenta as a mediator of stress effects on the developing brain^7-9^. Animal studies also have shown that the outcomes of altered placental functioning on neurodevelopment are substantially sex-specific, with males more vulnerable than females to prenatal adversity^8,9^. These observations raise the possibility that expression of schizophrenia risk associated genes may be different in placentae of male compared with female offspring and this might relate to the greater incidence of developmental disorders like schizophrenia among males^33,34^. We therefore tested whether PRS1 and PRS2 genes, which interact with ELCs on case-control status, are differentially expressed in placentae from male compared with female offspring. Analyses on control samples from the two datasets with sex information detected that the PRS1 and PRS2 genes are significantly enriched among the genes differentially expressed, and specifically up-regulated, in placentae from male compared with female offspring. (GSE35574: N=40 (17F, 23M), p=4.9e-08, Fig. 4a; GSE25906: N=37 (16F, 21M), p=2.3e-10; Fig. 4b). In the same datasets, the relative up-regulation was also present in male preeclamptic placentae (GSE35574: p=0.01; GSE25906: p=0.001). Analogous analyses in a heart dataset (GSE4172) and a fetal lung dataset (GSE14334) with sex information did not reveal up-regulation of the PRS1 and PRS2 genes in males compared with females (p>0.50). These data support a sex-biased role for the placenta in expressing genetic risk for schizophrenia.

**Fig. 4.**
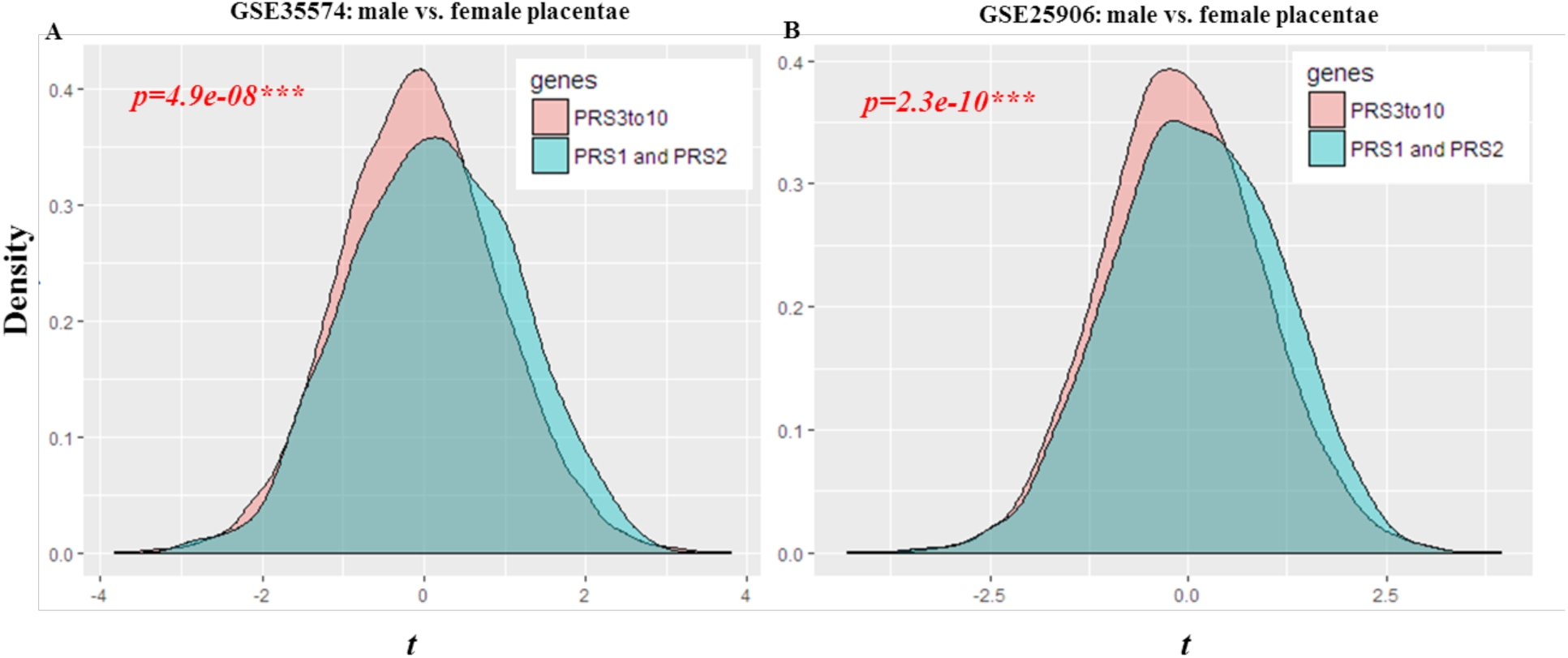
Schizophrenia risk genes are up-regulated in male compared with female placentae. **A-B).** PRS1 and PRS2 genes were tested for enrichment among the differentially expressed genes in placentae from male compared with female offspring. The figure shows the density plots of the T-statistics, from the differential expression analysis, of PRS1 and PRS2 genes (blue area), and of all the other genes in PRS’s (PRS3to10) constructed from lesser statistical thresholds (pink area). PRS1 and PRS2 genes are more likely to be differentially expressed and specifically up-regulated in placentae from male compared with female offspring (negative Tstatistics=more expressed in females; positive T-statistics=more expressed in males; ‘geneSetTest’ statistics at the top of the graphic).

## DISCUSSION

Here, we show that exposure to ELCs represents an early environmental context that influences cumulative genomic risk for schizophrenia derived from GWAS-significant loci. More to the point, the set of genes within these genomic loci that show interaction with intrauterine and perinatal complications is highly expressed in placenta, and the same set of genes displays differential enrichment in this tissue in abnormal invasive placental states. These results suggest that the most significant genetic variants detected by current GWAS^16^ contribute to risk of schizophrenia at least partly by converging on a developmental trajectory sensitive to intrauterine and perinatal adversity and linked with abnormal placentation. Moreover, the strikingly relative enrichment of expression of schizophrenia risk genes in placentae from male compared with female offspring suggests that gene-environment interactions influencing placental biology may account for the higher incidence of schizophrenia in males compared with females^33^.

Our results indicate a link between placental biology, ELCs and schizophrenia, even as the syndrome is diagnosed during early adult life, which resonates with a broader evolutionary perspective and the developmental trajectory of schizophrenia. Schizophrenia is thought to be a condition on which the human species has a monopoly, and the delayed emergence of the clinical disorder has been posited to reflect the relatively late maturation of highly evolved neural functions, such as prefrontal cortical circuitry^11^. Interestingly, the evolutionary complexity of the primate placenta shows parallels with the phylogenetically remarkable expansion of the human brain, particularly prefrontal cortical regions that are among the most affected in schizophrenia^35,36^; both placental complexity and brain expansion come with higher rates of ELCs in humans than in other species^35,37^. Our results are consistent with the possibility that some of the common genes implicated in schizophrenia risk – through diverse biological mechanisms – regulate the physiology of the placenta, the risk of ELCs, and thereby secondarily the development of the brain, potentially interacting with other mechanisms of gene regulation acting primarily within fetal brain^24,38^.

Despite many studies that have stressed the role of prenatal development and early life events in affecting risk for brain disorders like schizophrenia^5,12,13,39^, as well as autism^40,41^, the mechanisms by which this may happen have been elusive. Genetic research has been successfully focused on detecting GWAS-significant variants, but the difficulty of collecting environmental data has hindered defining the developmental context in which these common variants may have their critical effects. Our results underscore the importance of assessing early environmental factors such as obstetrical complications in addition to genetic risk, to fully investigate their joint effect on susceptibility to neurodevelopmental disorders. Our results also point to the placenta as a crucial mediator of this interaction in relation to schizophrenia in particular, but likely to other neurodevelopmental disorders in general, underscoring the need for further research on placenta physiology in the context of brain development and genomic risk. Pursuing this path should advance the role of prenatal care for reducing the burden of psychiatric illness and may identify new strategies^42^ for placental health as a form of primary prevention of schizophrenia, particularly in males with high genetic risk.

## METHODS

### Derivation of Polygenic Risk Profile Scores

Cumulative genetic risk profile scores (PRS’s) were calculated for each individual, as described elsewhere^16^. Briefly, PRS’s are a measure of genomic risk calculated as the weighted sum of risk alleles for schizophrenia from the recent GWAS study^16,20^. We thus multiplied the natural log of the odds ratio (OR) of each index SNP – from this recent schizophrenia GWAS^16^ - by the imputation probability for the corresponding reference allele at each variant, and summed the products over all variants, so that each subject had whole genome PRS’s as originally described for this measure and detailed in supplementary methods. For the *scz_lie_eur* sample, odds ratios (ORs) and 102K index SNPs were derived from a meta-analysis of PGC GWAS datasets excluding our discovery sample (PGC 2014, non-*scz_lie_eur* PGC GWAS)^16^. Consistently, for the *scz_munc_eur* and the *scz_gras_eur* samples (also included in the PGC GWAS^16^), ORs and index SNPs were derived from a meta-analysis of PGC GWAS datasets, excluding respectively the *scz_munc_eur* and the *scz_gras_eur* samples. For the *scz_bari_eur* and the *scz_osak_asi* samples, ORs and index SNPs were derived from the PGC GWAS datasets, since these samples are not included in the PGC GWAS dataset^16^. Consistent with the standard procedure for PRS calculation^16,20^, only autosomal SNPs were included in the analysis, in order to prevent any bias related to sex in the PRS calculation. We LD pruned and “clumped” the SNPs, discarding variants within 500 kb of, and in r2 ≥ 0.1 with, another (more significant) marker, as reported elsewhere^16^. Ten PRS’s (PRS1-PRS 10) were calculated using sub-sets of SNPs selected according to the GWAS p-value thresholds of association with schizophrenia: 5e-08 (PRS1), 1 e06 (PRS2), 1e-04 (PRS3), 0.001 (PRS4), 0.01 (PRS5), 0.05 (PRS6), 0.1 (PRS7), 0.2 (PRS8), 0.5 (PRS9), and 1 (PRS10). SNPs in sets with lower p-values are also in sets with higher p-values (e.g. SNPs in PRS1 are included in PRS2, SNPs in PRS2 are included in PRS3, and so on). A detailed list of SNPs included in PRS1 and PRS2 is provided in Table S2. We performed all the analysis both including and excluding the top GWAS significant SNP in the extended MHC locus (chr6:25-34 Mb), with similar results (Table S4). For additional analyses (Figure S21), we also calculated PRS’s from sets of SNPs with higher p-values (PRS3to10) without including SNPs in sets with the lowest p-values (PRS1 and PRS2). Details for genotyping and PRS calculation of each sample are reported in SI.

### Assessment of Early Life Complications (ELCs)

ELCs are here referred to as “somatic complications and conditions occurring during pregnancy, labor-delivery and the neonatal period” potentially harmful for the offspring, with special focus on the central nervous system (CNS)^21^. These events are also referred to elsewhere as ‘obstetric complications’^14,21^ and, despite their potential frequent occurence^43^, do not lead to negative outcomes in most cases. We assessed ELCs through medical records, when available, and personal interviews, described as follows:

*scz_lie_eur*, and *scz_bari_eur*: we used mainly maternal recall based on an extensive personal interview, which has been repeatedly shown to represent a reliable method for obtaining ELCs history, when used in a careful and detailed manner^44,45^. Specifically, we used a well-standardized and validated questionnaire^14^, based on all the items scored with the McNeil–Sjöström scale for obstetric complications^21^. It covers the entire duration of pregnancy and early neonatal life, and also contains indicators of reliability, and an assessment of the seriousness of each complication (see SI for details);
*scz_osak_asi*: we used mainly medical records. When medical records were not exhaustive, we interviewed the mothers of the patients; the histories were again scored directly based on the McNeil-Sjöström metrics;
*scz_munc_eur* and *scz_gras_eur*: we used medical records, including all the discharge letters of patients, and personal interviews administered to the patients. Differently from the questionnaires used in the other samples, these interviews did not contain all the items included in the McNeil–Sjöström scale, thus increasing the risk of incomplete information. History of ELCs from the available information was again scored using the McNeil–Sjöström scale^21^.

In the McNeil–Sjöström scale^21^ each ELC is assigned a severity level of 1-6, based on the degree of inferred potential harm to the offspring CNS. ELCs with severity weight ≥ 4 are considered potentially clearly harmful or relevant factors in fetal stress. The McNeil– Sjöström scale in the context of maternal recollection has been well validated in comparison to hospital records^21^. As in other reports^21,46,47^, we defined a positive history of ELCs based on the presence of at least one serious ELC (severity level ≥4), and we identified the severity level of each individual as the highest severity level of all the ELCs occurring in that individual. GWAS derived PRS’s were unknown to both the individuals who provided the information about ELCs and to the researchers who collected and evaluated them. Individuals were excluded if the information provided was incomplete or inconsistent (e.g. contradictory answers to related questions), or if the presence of a complication was certain but a severity weight could not be confidently determined. The frequency of ELCs in our samples may be not representative of the general populations (see Supplementary Information: “Considerations about the assessment and frequency of ELCs”). Table S20 contains a list of the ELCs detected in each sample.

### Selection of PRS1 and PRS2 genes

In order to define genes mapping to the PRS1 and PRS2 loci for gene set analyses, we used two alternative criteria:

PGC LD regions: we considered, as PRS1 and PRS2 genes, all the UCSC genes overlapping the LD-regions associated with each SNP (R^2^>0.6), as reported in a previous reference^16^ and on the PGC website <u>(http://www.med.unc.edu/pgc/downloads);</u>
distance: we considered, as PRS1 and PRS2 genes, all the UCSC genes mapping 500kb +/- the index SNPs of each PRS in the discovery sample (*scz_lie_eur*). We use this criterion, in addition to the ‘traditional’ LD criterion, on the grounds that LD differs among populations, as we analyzed multiple samples. Moreover, the LD regions associated with each SNP have a huge variability: for example, two out of the 108 GWAS-significant schizophrenia-risk SNPs have an LD-region that spans only 1 bp (rs4766428, rs117074560)^16^. Further, it has been shown that GWAS SNPs are often associated with expression of genes that are not their nearest genes and are outside the associated LD-regions^16,48^. Finally, the distance of 500kb +/- the index SNPs is within the range commonly used for detection of cis-eQTLs^48^ and is the same dimension used to calculate PGC loci eQTLs in the original PGC report^16^. This criterion allowed us to distribute 21028 out of 23056 UCSC genes among the 10 PRS’s.

Differences between the two list of genes (reported in Table S9a-b), are related not only to the criterion adopted for SNP selection (distance or LD), but also to the fact that the PGC loci associated with schizophrenia at p<5-08 are defined based on combining the primary GWAS and the supplementary deCODE data, while SNPs for PRS calculation are derived from the primary GWAS only^16^. We excluded from our analysis genes that were irrelevant to our question, i.e. mitochondrial, X and Y-chromosome genes or other genes mapping to loci not used for PRS’s calculation, thus avoiding the risk of overinflating p-values <u>(https://bioconductor.org/packages/release/bioc/vignettes/GOstats/inst/doc/GOstatsHyperG.pdf)</u>. After exclusion of the genes on sex chromosomes and genes undetected in the expression datasets analyzed, the final number of PRS1 and PRS2 genes was 1643 in the list based on distance (matching 325 out of the 348 genes assigned to the 108 schizophrenia GWAS significant loci^16^), and 589 in the gene list based on LD (matching 334 out of the 348 genes assigned to the 108 schizophrenia GWAS significant loci^16^). In both gene lists, PRS1 genes are a subset of PRS2 genes (therefore referred in the text as PRS1 and PRS2 genes). We performed all the gene set analyses, with PRS1 and 2 genes defined with both criteria (LD and distance), and found consistent results (Table S9c). In the main text, we report results with the PRS1 and PRS2 genes defined based on the distance criterion (Table 2).

## Acknowledgments

We are grateful to the Lieber and Maltz families for their visionary support that funded the analytic work of this project. We thank all the participants in the study and their families. We thank Joo Heon Shin, for help with the gene expression analysis, Marina Mancini, Raffaella Romano, Rita Masellis, for help with data acquisition. We also thank Sally Cheung and John Meyer for data management, Susan Fisher for guidance with placental datasets and review of the manuscript, and Brion Maher and Michael Neale for their thoughtful reviews of the data and manuscript. We thank the Psychiatry Genomics Consortium for providing the statistics for PRS calculation. We also thank all the authors of the publicly available placental datasets that have been used in this study. The collection of the ELC and genetic data for the American samples was supported by direct funding from the Intramural Research Program of the NIMH to the Clinical Brain Disorders Branch (DR Weinberger, PI, protocol 95-M-0150, NCT00001486, annual report number: ZIA MH002942-03 CTNB). The GRAS data collection has been supported by the Max Planck Society, the Max Planck Förderstiftung, the DFG (CNMPB), EXTRABRAIN EU-FP7, and the Niedersachsen-Research Network on Neuroinfectiology (N-RENNT).

## Contributions

G.U., G.P. and D.R.W. designed the study and interpreted the results. G.U., G.P., J.F.R., E.G.H., A.E.J. and C.C. carried out statistical analyses. G.U., Q.C., M.M., R.E.S., H.E. and D.R.W. organized and performed genotyping, imputation and risk profile scoring. G.U., S.M., M.B., J.S., K.F.B., M.F.E., R.E.S., G.B., R.H., D.R., H.E., A.B. and D.R.W. organized and carried out subject recruitment and biological material collection in the discovery sample and in the replication samples, while G.U., G.P., S.M., A.P, G.M., M.B., H.Y., R.H., D.R. and H.E. carried out early life complications assessment. J.F.R. and E.G.H. contributed to the collection of the placental tissue used in the RNAseq analysis and, together with G.U., G.P., C.C. and D.R.W. interpreted the results of the gene set enrichment analysis in placental samples from complicated pregnancies compared with controls. G.U., G.P. and D.R.W. drafted the manuscript and all authors contributed to the final version of the paper.

## Supplementary Information

Supplementary Materials and Methods

Supplementary Table, Figures and Results:

Tables S1, S3-8, S10-13, S18-20

Figures S2-5, S7-21(Fig. S1 and S6 in separate files)

Captions for additional Tables S2, S9, S14-17

Supplementary References (1-47)

Additional Tables S2, S9, S14-17

